# Promoter DNA Methylation Epigenetically Regulates the Tumor-Suppressor Function of the miR-379/656 (C14MC) Cluster in Hepatocellular Carcinoma

**DOI:** 10.64898/2025.12.02.691781

**Authors:** Shreyas Hulusemane Karunakara, Shama Prasada Kabekkodu, Akila Prashant, Rohit Mehtani, Prashant Vishwanath, Gopalakrishna Ramaswamy, Prasanna Kumar Santhekadur

**Author notes:** **Corresponding author’s**. **Email ID of coauthors** Shreyas Hulusemane Karunakara, Shama Prasada Kabekkodu, Akila Prashant, Rohit Mehtani, Prashanth M Vishwanath, Gopalakrishna Ramaswamy.

## Abstract

Hepatocellular carcinoma (HCC) is a leading cause of cancer-related mortality. Several microRNAs (miRNAs) play key roles in HCC development and progression. The role of epigenetic processes like DNA methylation in the regulation of miRNAs is critical to HCC pathogenesis. In this study, we show that the miR-379/656 cluster (C14MC) acts as a tumor suppressor cluster and is epigenetically regulated by DNA methylation.

We demonstrated that C14MC is downregulated in HCC cell lines through nCounter assay and in clinical samples from The Cancer Genome Atlas (TCGA) liver hepatocellular carcinoma tissues. The C14MC promoter was identified and characterized through cloning and dual luciferase assay. Furthermore, we demonstrated that the loss of C14MC tumor suppressor function is directly regulated by the hypermethylation of promoter-bound CpGs, as shown in artificial methylation experiments. The reactivation of specific C14MC miRNAs, like miR-299-5p and miR-376c-3p via mimics, abrogated the expression of several target oncogenes, including PARP1, SPP1, RAD21, and CENPA that regulate critical molecular pathways such as the p53 signaling and NF-kappa B signaling pathways in HCC. Additionally, overexpression of miR-299-5p and miR-376c-3p inhibited HCC cell migration and invasion, suggesting that overexpression of candidate C14MC miRNAs can mitigate cancer hallmarks in HCC cells. Furthermore, we checked for the clinical correlation of C14MC targets and their target genes in terms of survival outcomes and identified the key genes associated with prognostic potential in HCC.

We conclude from our experimental findings that C14MC is a tumor-suppressor miRNA cluster and is regulated epigenetically by methylation in HCC. Several of these miRNAs and their targets can be used for early HCC diagnosis and prognosis. Thus, targeting C14MC can be useful in HCC management.

## 1. Introduction

Hepatocellular carcinoma (HCC) is the most commonly reported form of primary liver cancer. In 2021, HCC added around 529,000 cases and 484,000 mortalities to the existing cancer burden reported globally (1). While hepatitis B remains the leading factor, steatohepatitis is an emerging contributor of HCC (2). While the existing vaccination programs against HBV have reduced the HBV-associated HCC burden, the lack of a clear understanding of critical molecular players in pathogenesis has resulted in poor diagnosis and prevention of the disease at an advanced stage, thereby increasing the incidence and mortality associated with HCC (3). Thus, there is a need to understand how various genetic and epigenetic factors operate during HCC pathogenesis and progression to identify reliable markers for clinical management of the disease.

Non-coding RNAs (ncRNAs) are a broad class of regulatory transcripts that perform critical functions associated with gene expression regulation (4). These RNAs operate at transcriptional and post-transcriptional levels through various mechanisms like gene silencing, epigenetic regulation, such as DNA methylation and chromatin structure remodeling, and histone modifications (5,6). The most widely studied non-coding RNAs comprise a class of small-ncRNAs called microRNAs (miRNAs) with potential implications in disease pathogenesis, including cancers (7). Aberrant expressions of miRNAs can cause either gain or loss of function effects depending upon the site of expression and type of cancer (8–10). Additionally, dysregulated miRNA expression might affect the downstream transcriptome and promote pathways and processes involved in cancer development and progression (11–12). We and many others have previously reported that aberrant miRNA expressions can promote HCC development (13–16). Therefore, quantifying the miRNA expression profiles can be of diagnostic and prognostic potential for HCC. Interestingly, several miRNAs are coexpressed and coregulated in unison as clusters (17–20), and the role of epigenetic processes like DNA methylation operating at critical regulatory genomic locations such as the promoters and enhancers of these clusters at large, might help understand the intricate regulatory processes that affect cluster expressions and their biological processes and downstream pathways by regulating their target transcriptomes (21–22).

The miR-379/656 cluster, also known as the chromosome-14 miRNA cluster (C14MC), is the second-largest miRNA cluster. Located on the 14q32.31 locus, alongside other imprinted coding (DLK1, DIO3, and RTL1), non-coding genes (MEG3 and MEG8), and small nucleolar RNAs, this cluster encodes around 50 different miRNAs that are regulated by a single promoter region (23–24). The expression of C14MC is mainly reported in tissues of epithelial origin (25). The aberrant C14MC expression is associated with multiple diseases and developmental disorders (26). The abnormal expressions of several C14MC members are implicated as either oncogenic or tumor suppressor in several cancers (23, 27–29). In addition, studies have shown that promoter-bound alterations or changes in other internal regulators of the C14MC might be the underlying cause for disruption in C14MC expression (30).

Several studies have shown miRNAs of this cluster to be aberrantly expressed in HCC. Members like miR-656, miR-409, and miR-379 are downregulated in HCC. Specifically, miR-656 downregulation is known to promote HCC proliferation via SIRT5 upregulation. Further miR-656 upregulation inhibited the invasion and migratory potential of HCC cells (31). Another miRNA of the cluster, miR-379, which was reported to be underexpressed in HCC, was also shown to be associated with TNM stage and metastasis. Additionally, an ectopic expression of this miRNA suppressed HCC migration, invasion, and EMT through targeting the FAK/AKT signaling (32). Another study pointed towards enhanced HCC invasion and metastasis due to the tumor suppressor function of miR-409 of C14MC, which resulted in BRF2 overexpression and Wnt/β-Catenin activation (33). However, the expression of the complete cluster in HCC and the role of epigenetic mechanisms like DNA methylation in regulating the cluster are not understood. In this study, we have demonstrated that the C14MC acts as a tumor suppressor cluster and that promoter hypermethylation can be a key epigenetic driver of this downregulation. Further, we have predicted various oncogene targets of the cluster, including PARP1, SPP1, CENPA, and RAD21, whose expressions are regulated by C14MC miRNAs. Furthermore, restoring C14MC miRNAs like miR-299-5p and miR-376c-3p inhibited HCC cell migration and invasion. Additionally, we have also evaluated the diagnostic and prognostic potential of C14MC and its target genes that can be used as biomarkers for HCC.

## 2. Results

### 2.1. HCC exhibits miR-379/656 cluster downregulation

The analysis of nanostring nCounter data showed downregulation of different miRNAs belonging to C14MC. We observed that cluster miRNAs like miR-376c-3p and miR-382-5p were downregulated in HepG2 cells (**Figure 1A-1B**). Other cluster miRNAs, miR-299-5p, miR-376c-3p, miR-656-3p, miR-376a-3p and miR-377-3p were downregulated in Huh7 cells (**Figure 1C-1G**). We could not detect other C14MC miRNAs to be significantly downregulated in HCC cells. However, the TCGA-LIHC data confirmed the downregulation of around 30 C14MC miRNAs in primary HCC tissues compared to normal liver tissues (**Figure 1H**). Further, the combined sensitivity and specificity associations revealed an area under the curve (AUROC) of 0.8890 (95% CI, p<0.0001) (**Figure 1I**) for miRs-376c-3p and −382-5p, and an AUROC of 0.8067 (95% CI, p<0.0001) for miRs-299-5p, miR-376c-3p, miR-656-3p, miR-376a-3p and miR-377-3p (**Figure 1J**), respectively.

**Figure 1.**
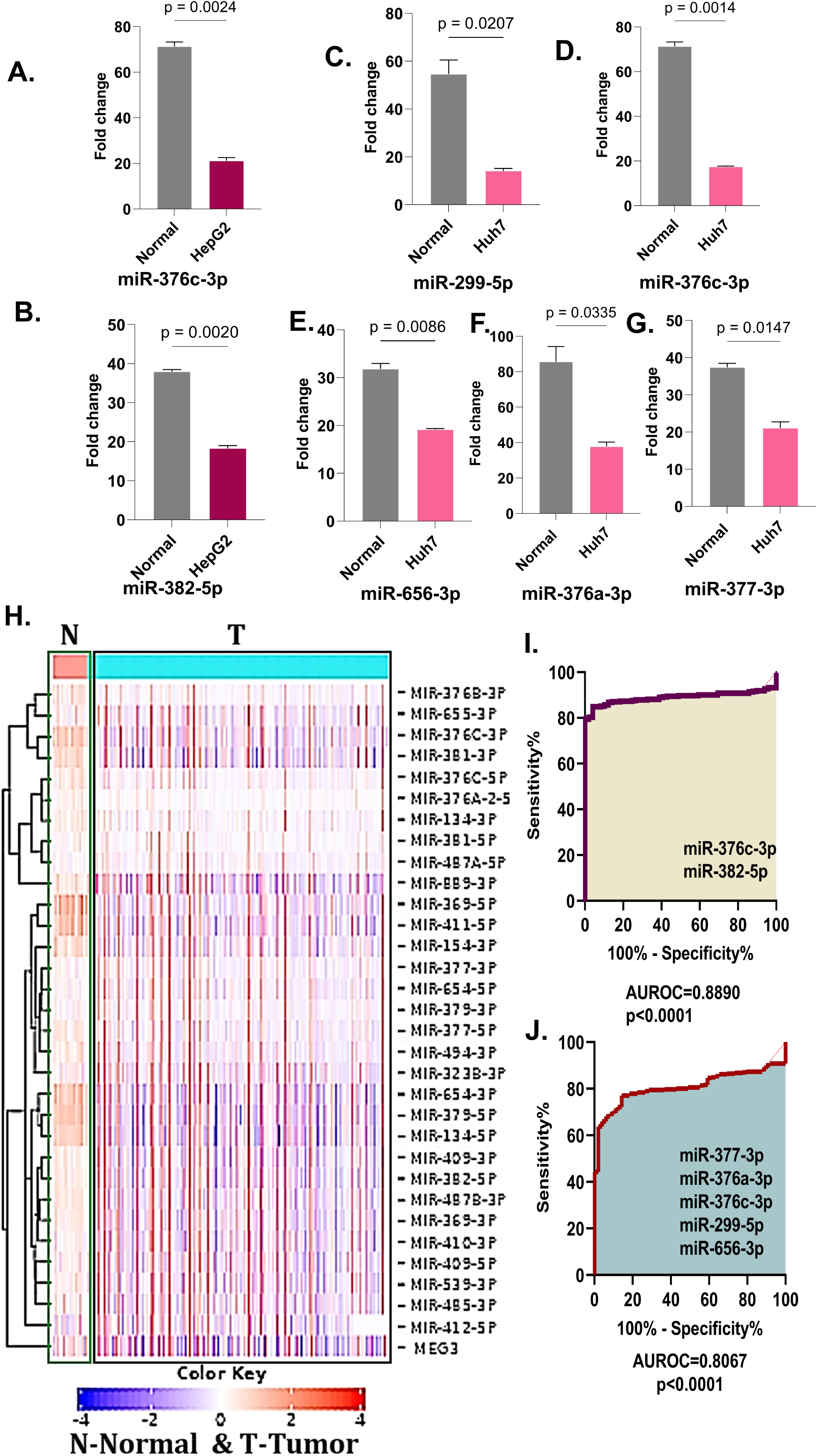
C14MC expression analysis in HCC. (A)-(G) Bar graphs showing differential downregulation of C14MC member miRNAs in HepG2 and Huh7 HCC cell lines compared to control RNA. (H) Heatmap showing differential downregulation of DLK1-DIO3 gene MEG3 and around C14MC miRNAs in primary HCC samples in comparison with adjacent normal liver tissues from the TCGA-LIHC cohort. (I) and (J) ROC curves for C14MC were significantly downregulated in HepG2 and Huh7 cell lines using TCGA-LIHC expression data. The sum of the expressions of respective C14MC miRNAs is highlighted. The error bar represents the mean ± SD of duplicate experiments. Statistical significance was determined by an unpaired Student’s *t*-test.

### 2.2. Characterization of miR-379/656 (C14MC) cluster promoter

We predicted the possible promoter region of C14MC relative to its transcription start site (TSS) using the FANTOM5 tool. The annotated promoter, with coordinates (chr14:101,291,001–101,292,434) located within the MEG3 region. The region exhibited several promoter-associated markers, including multiple cis-regulatory elements, a DNase I hypersensitivity region, H3K27Ac, and a CpG island (**Figure 2A**). The associations of these genomic signatures suggested that the predicted region likely functions as the probable putative promoter of the cluster. The interactive map of the promoter region is shown in **Figure 2B**. The successful cloning of the promoter was confirmed by isolating the plasmids from transformed *E. coli-*DH5α cells and subsequently performing double and diagnostic digestions (**Figure 2C-2E**). We then confirmed the promoter function of this 1,434 bp region by dual luciferase assay. We observed a nearly 2-fold increase in luciferase activity in pB-C14MC containing the promoter in comparison with the pGL3-Basic (**Figure 2F**). We then assessed the effect of DNA methylation on the promoter activity by artificially methylating the pB-C14MC full-length construct and measuring the luciferase activity. The artificial DNA methylation showed a significant reduction in the promoter activity compared to the unmethylated construct (**Figure 2G**). Additionally, we identified different transcription factors (TFs) that might bind to the C14MC promoter region using the TRASFAC (https://genexplain.com). We identified that HNF4α, MEF2A, PBX3, c-Myc, NGFIβ, TBP, SMAD4, KLF4, PRDM1, TBX5, SOX6, TCF-7L2, NF-Y, and CEBPα binding sites were present within the C14MC promoter (**Figure 2H**). DNA methyltransferases, primarily *de novo* (DNMT3A, DNMT3B, DNMT3L) and maintenance methyltransferases (DNMT1, DNMT2), are the enzymes that catalyse DNA cytosine methylations (34). We have previously reported from TCGA-LIHC data that DNMT1, DNMT3A, and DNMT3B are upregulated in HCC (16). These experiments suggest that the C14MC upstream region identified to be likely the promoter region, whose function is regulated by methylation levels.

**Figure 2.**
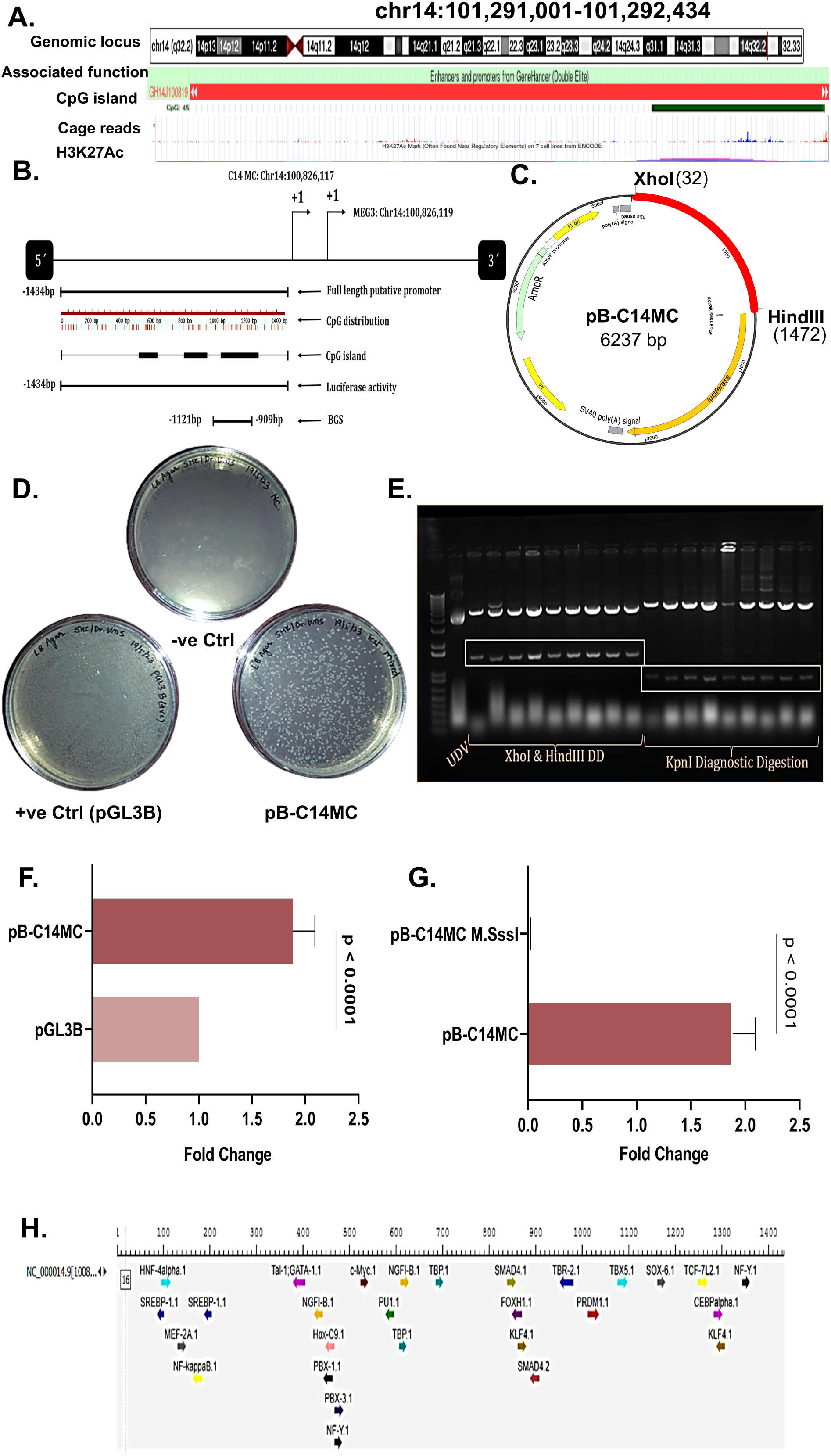
Identification and characterization of the C14MC promoter. (A)The map of the putative promoter of C14MC showing CAGE reads, CpG islands, and histone acetylation marks. (B) Interactive map showing promoter CpG distribution and regions of promoter characterized using methylation studies and luciferase assay-based characterization. (C) Vector map of recombinant clone pB-C14MC. (D) LB agar plates showing transformed E.coli DH5α colonies with recombinant pB-C14MC. (E) Agarose gel electrophoresis results confirming the successful cloning of the full-length promoter by XhoI and HindIII double digestion (1434 bp) and orientation confirmation by KpnI digestion (801bp). The reference ladder used is a 100 bp ladder. (F) The relative fold change in luciferase activity, showing a 2-fold increase in HepG2 cells transfected with pB-C14MC, showing promoter activity associated with the cloned region. (G) Artificial methylation experiment results showing significant reduction in promoter function upon M.SssI methyltransferase treatment and subsequent transfection to HepG2 cells, followed by DLR assay. (H) The possible transcription factors that can bind to the C14MC promoter in both forward and reverse genomic orientations.

### 2.3. HCC exhibits promoter-specific hypermethylation patterns

We further investigated the associations of the exact methylation status of promoter-bound CpGs and correlated the methylation levels with the C14MC expressions. A bisulfite PCR and sequencing targeting a 212bp (−1121 to −909 bp) promoter region were performed to detect the partial promoter methylation status (**Figure 3A-3B**). We observed that CpG-12 showed relatively lower methylation (1-10%) in both HepG2 and Huh7 cells, while other CpGs-8, 9, 10, 13, 14, 15, 16, 17, and 18 showed hypermethylation (90-100%) in HepG2 cells, and CpGs-10, 11, 14 and 17 were significantly hypermethylated in Huh7 cells (**Figure 3C**). In the TCGA-LIHC dataset, we identified that C14MC promoter CpGs-cg14245102, cg26374305, and cg04291079 were hypermethylated compared to adjacent normal tissues (**Figure 3D**). However, we could not find any stage-wise associations of methylation status of these three CpGs in LIHC (**Figure 3E**). We further validated the hypermethylation status of the three promoter CpGs by validating in an independent cohort from the Genome Expression Omnibus (GEO): GSE54503, containing paired normal and HCC samples, where CpGs (cg14245102, cg26374305, cg04291079) were hypermethylated in paired HCC tissues (**Figure 3F**). We also observed in another independent cohort (GSE78732), the C14MC associated CpGs-cg10065153, cg10943497, cg14034270, cg02888166, cg15419911, cg11035687, and cg14121301 as hypermethylated in HCC. Additionally, it was seen that CpGs relative to the position of the island, that is, the N-shore and the S-shelf, were relatively hypermethylated, while the island-bound CpGs had relatively stable methylation levels in HCC tissues compared with normal tissues (**Figure 3F**).

**Figure 3.**
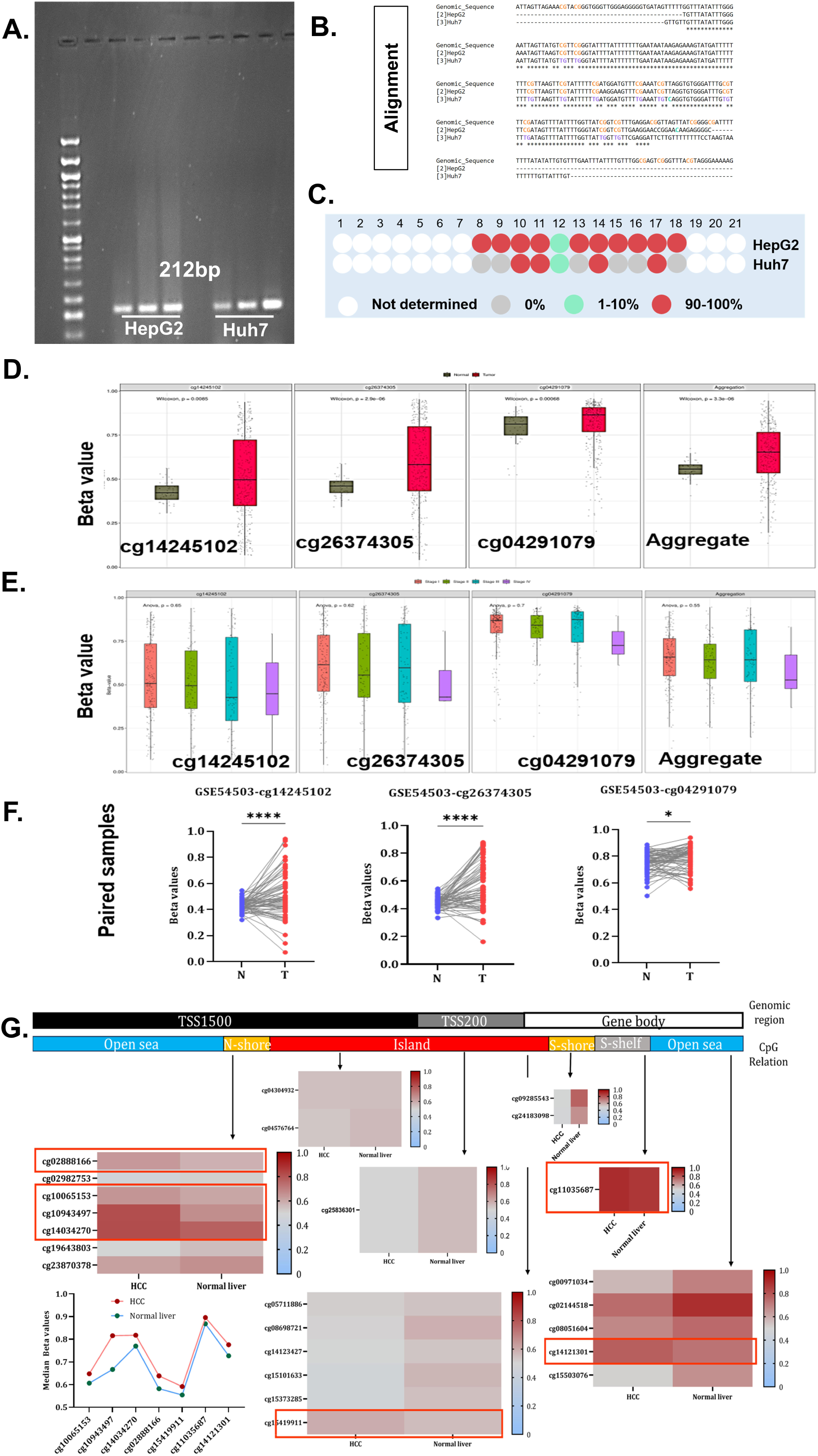
DNA hypermethylation regulates C14MC expression in HCC cells and tissues. (A) Agarose electrophoresis image under UV showing C14MC bisulfite PCR amplicons of 212bp from HepG2 and Huh7 cells in biological triplicate. (B) Multiple sequence alignment results showing alignment of reads obtained from Sanger sequencing of a 212bp bis-PCR product against the reference promoter sequence. The CpG alignments within the promoter region are highlighted in orange/ purple colors, respectively. (C) Differential methylation of different CpG sites exhibiting hypermethylation in HCC cells. (D) The differential methylation data from the TCGA-LIHC cohort show significant hypermethylation of promoter CpGs-cg14245102, cg26274305, and cg04291079. (E) Stage-wise methylation analysis of CpGs-cg14245102, cg26274305, and cg04291079 in the TCGA-LIHC cohort, which showed no significant methylation changes. (F) Results validating differential hypermethylation of cg14245102, cg26274305, and cg04291079 CpGs in GSE5450, containing 10 paired normal and HCC tissues. (G) Differential heat-map mapping different C14MC promoter CpGs to genomic regions, including island, shore, shelf, and sea regions from EWAS datahub. Specific C14MC promoter-bound CpGs exhibiting a median increase in hypermethylation mapped to these regions are highlighted.

### 2.4. Identification and validation of C14MC cluster target interactome in HepG2 cells

We identified the targets of downregulated C14MC candidate miRNAs in HepG2 cells using an *in silico* approach. Further, these targets were validated in HepG2 cancer spheroids by RNA-Seq. The target prediction pipeline included only the genes that had an inverse correlation with individual C14MC miRNA expression, or only the target genes that were upregulated in HCC were considered for expression profiling. We observed that in HepG2 spheroids, miR-376c-3p targets; ANK1, ATAT1, ELF4, CENPA, and CHEK2 to be the target oncogenes (**Figure 4A-4C**). The ontologies associated with these genes included biological processes (BP) such as DNA damage induced protein phosphorylation, NK-T cell proliferation, establishment of cell polarity, cytokinesis, microtubule organization; molecular functions (MF): insulin-activated receptor activity, tubulin N-acetyltransferase activity, cytoskeletal anchor activity, G-protein alpha subunit binding, PI3-Kbinding and protein serine/threonine/tyrosine kinase activity; and pathways: Insulin/IGF mediated MAPK cascade, p53 signaling, and PDGF signalling pathways (**Figure 4D**). Likewise, the miR-382-5p target genes-SERGEF, MTHDF2, SYNJ, UBB, DAD1, TM4SF, ATG10, NOL4L, SLC6A8, RPLP0, YBX1, and TRPV2 (**Figure 4E-4F**) were found to be enriched in critical MFs such as methenyltetrahydrofolate cyclohydrolase/ dehydrogenase activity, phosphate ion binding, phosphatase activity; and the key pathways identified were formyltetrahydrofolate biosynthesis, ACE2/ ACE4 signaling, and Nicotinic acetylcholine receptor signaling cascades (**Figure 4G**).

**Figure 4.**
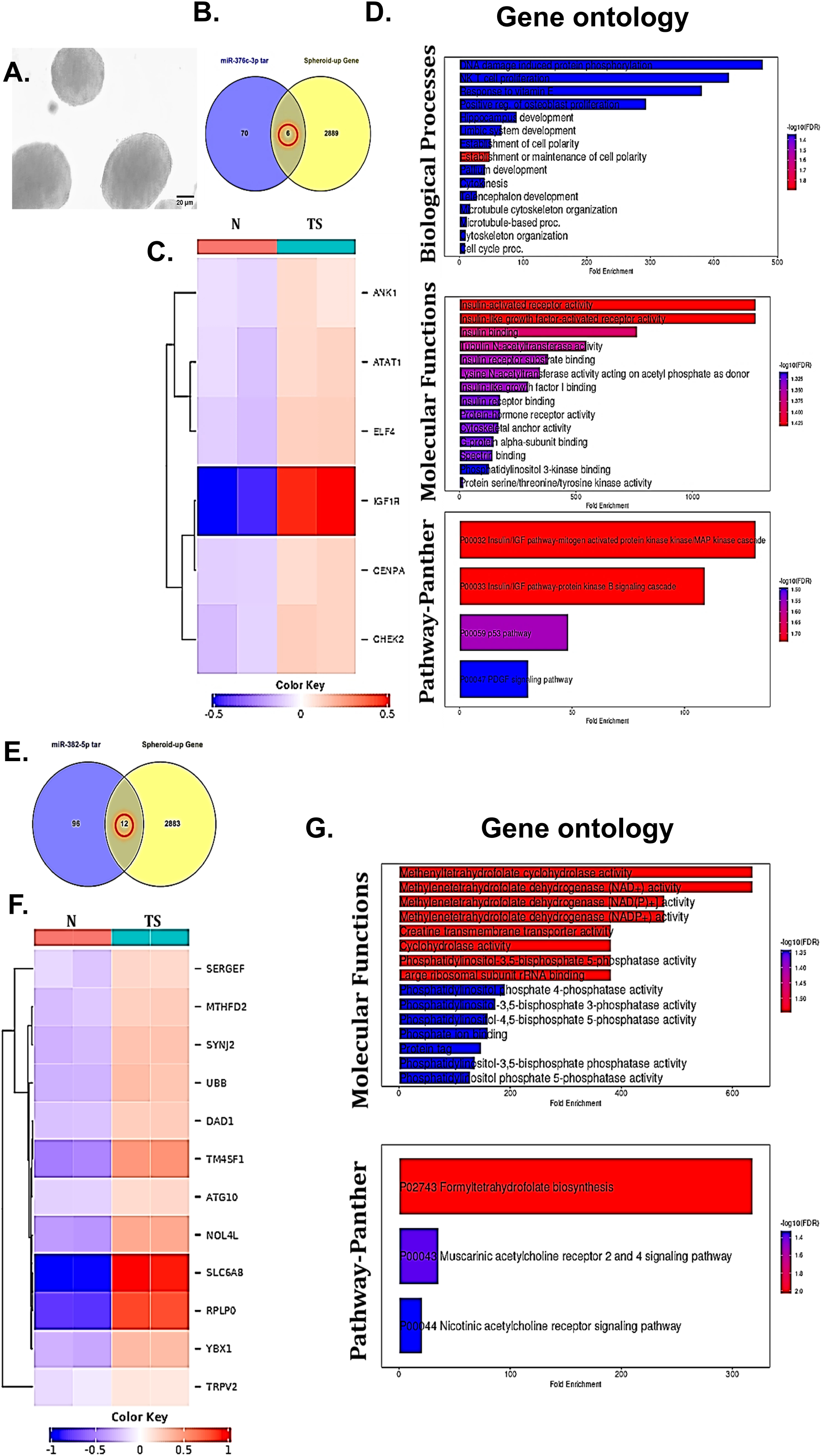
Validation of C14MC target gene expressions in HepG2 spheroids. (A) Micrographs showing HepG2 spheroids developed using a modified force-suspension method. (B) and (C) Transcriptomic analysis showing the expressions of 6 differentially overexpressed gene targets of miR-376c-3p (D) Gene ontology analysis showing significant biological processes, molecular functions, and pathways regulated by the miR-376c-3p target genes. (E) and (F) Transcriptomic analysis showing the expressions of 12 differentially overexpressed gene targets of miR-382-5p (D) Gene ontology analysis showing significant molecular functions, and pathways associated with these 12 miR-382-5p target genes.

### 2.5. Overexpression of miR-379/656 cluster miRNAs abrogates the expression of target oncogenes in Huh7 cells

The post-transcriptional network associated with C14MC targets in Huh7 cells was identified using an *in silico* approach and further validated by overexpressing specific miRNAs and measuring the target gene expression by qRT-PCR. We identified SPP1, IGFBP3, RAD21, CENPA, PARP1, LMNB1, SERPINE1, ESR1, CXCL2, and CDT1 as top ten hub genes of candidate C14MC (miR-299-5p, miR-376c-3p, miR-376a-3p, miR-656-3p, and miR-377-3p) (**Figure 5A**), which regulated critical gene ontologies such as chromosome organization, protein localization, and cell communication regulation (BP) (**Figure 5B**) and p53 signaling, NFκB signaling, apoptosis, apelin signaling and Cellular senescence were the key enriched pathways (**Figure 5C**). Moreover, we could identify that genes PARP1, RAD21, SPP1 (miR-299-5p targets), and CENPA (miR-376c-3p target) were directly associated with liver inflammation and HCC (**Figure 5D**) and were upregulated in HCC tissues than normal in TCGA-LIHC (**Figure 5E-5H**). Hence, we decided to profile the expressions of these targets to check if their expressions are mitigated upon miRNA overexpression. We observed a significant reduction in the mRNA expression levels of all four oncogenes-PARP1, SPP1, RAD21, and CENPA upon overexpressing miR-299-5p or miR-376c-33p in Huh7 cells compared to the cells transfected with negative control (**Figure 5I-5L**), suggesting the potential reversal of target oncogene expression upon restoring component C14MC miRNAs.

**Figure 5.**
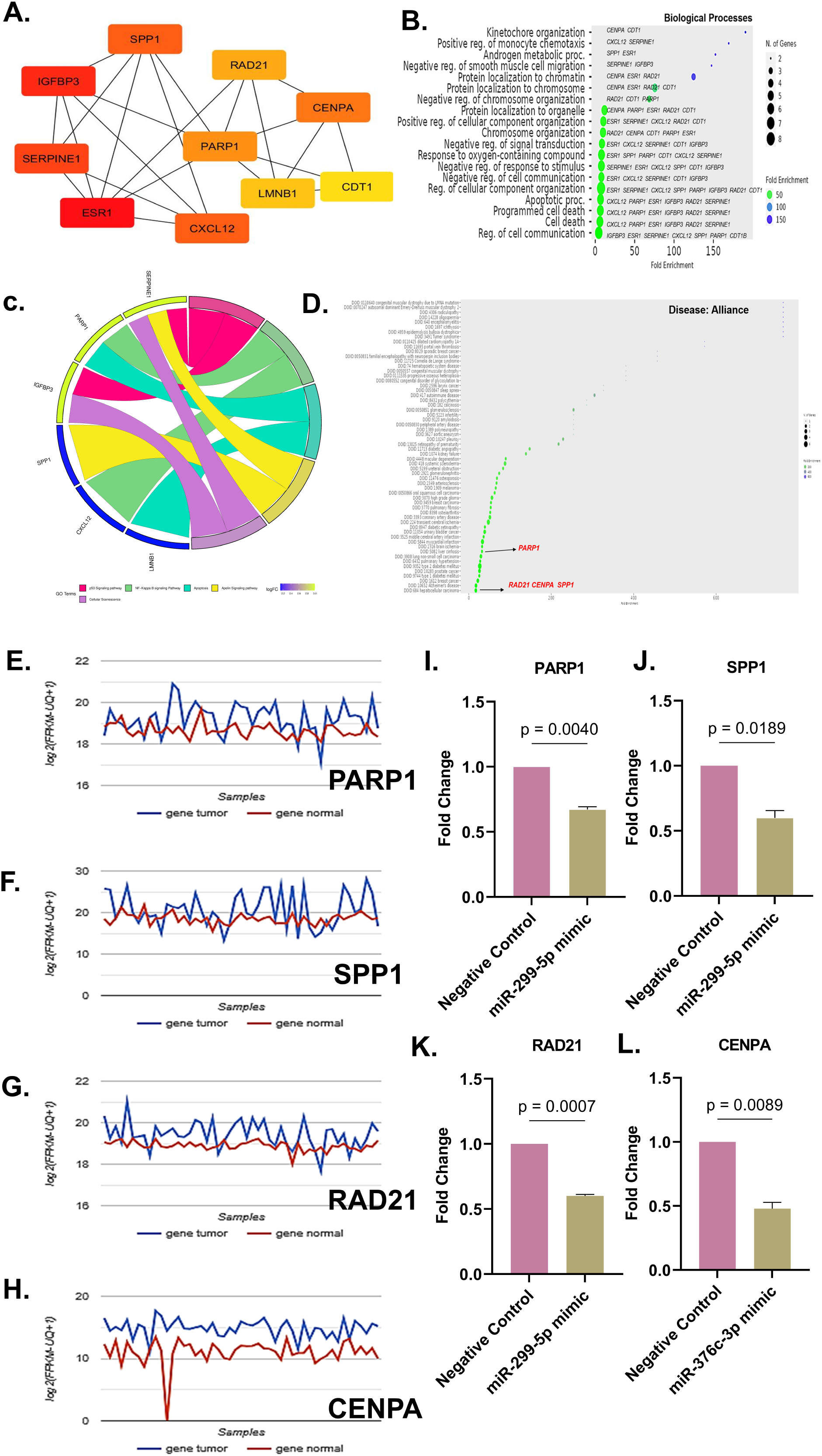
Restoring C14MC miRNAs mitigates the expression of target oncogenes in Huh7 cells. (A) Hub genes targeted by miR-299-5p, miR-376c-3p, miR-376a-3p, miR-377-3p, and miR-656-3p in Huh7 cells. (B) Enrichment analysis of hub genes showing the top biological processes-kinetochore organization, chromosome organization, and cell death. (C) Chord diagram showing key pathways regulated by C14MC hub genes. (D) Disease associations showing C14MC hub genes enriched in different diseases. PARP1, RAD21, CENPA, and SPP1 were enriched in liver inflammation and HCC. (E)-(H) Differential expression of C14MC target transcripts-PARP1, SPP1, RAD21, and CENPA in TCGA-LIHC showed a significant upregulation in HCC tissues (blue) in comparison to adjacent normal liver tissues (red). (I)-(K) Bar graphs showing fold change in expressions upon transfecting miRNA mimics to Huh7 cells. Ectopic expression of C14MC miRNA-miR-299-5p reduced the expression of target genes-PARP1, SPP1, and RAD21, and (L) Ectopic expression of C14MC miRNA-miR-376c-3p reduced the expression of CENPA, compared to cells transfected with negative control. Statistical significance was determined by an unpaired Student’s *t*-test.

### 2.6. Overexpressing of miR-299-5p and miR-376c-3p mimics inhibit the migration and invasion of Huh7 cells

We investigated the effect of C14MC expression restoration on cell migration and invasion, which are prominent hallmarks directly linked to HCC progression and metastasis. We observed that Huh7 cells transfected with either miR-299-5p or miR-376c-3p mimics showed reduced cellular migration (**Figure 6A-6C**) compared to the cells transfected with negative control. Likewise, overexpression of C14MC miRNAs-miR-299-5p and miR-376c-3p inhibited the invasion of Huh7 cells into the soft-agar spots (**Figure 6D-6F**). These experiments suggest that restoring C14MC expression might mitigate HCC hallmarks such as migration and invasion.

**Figure 6.**
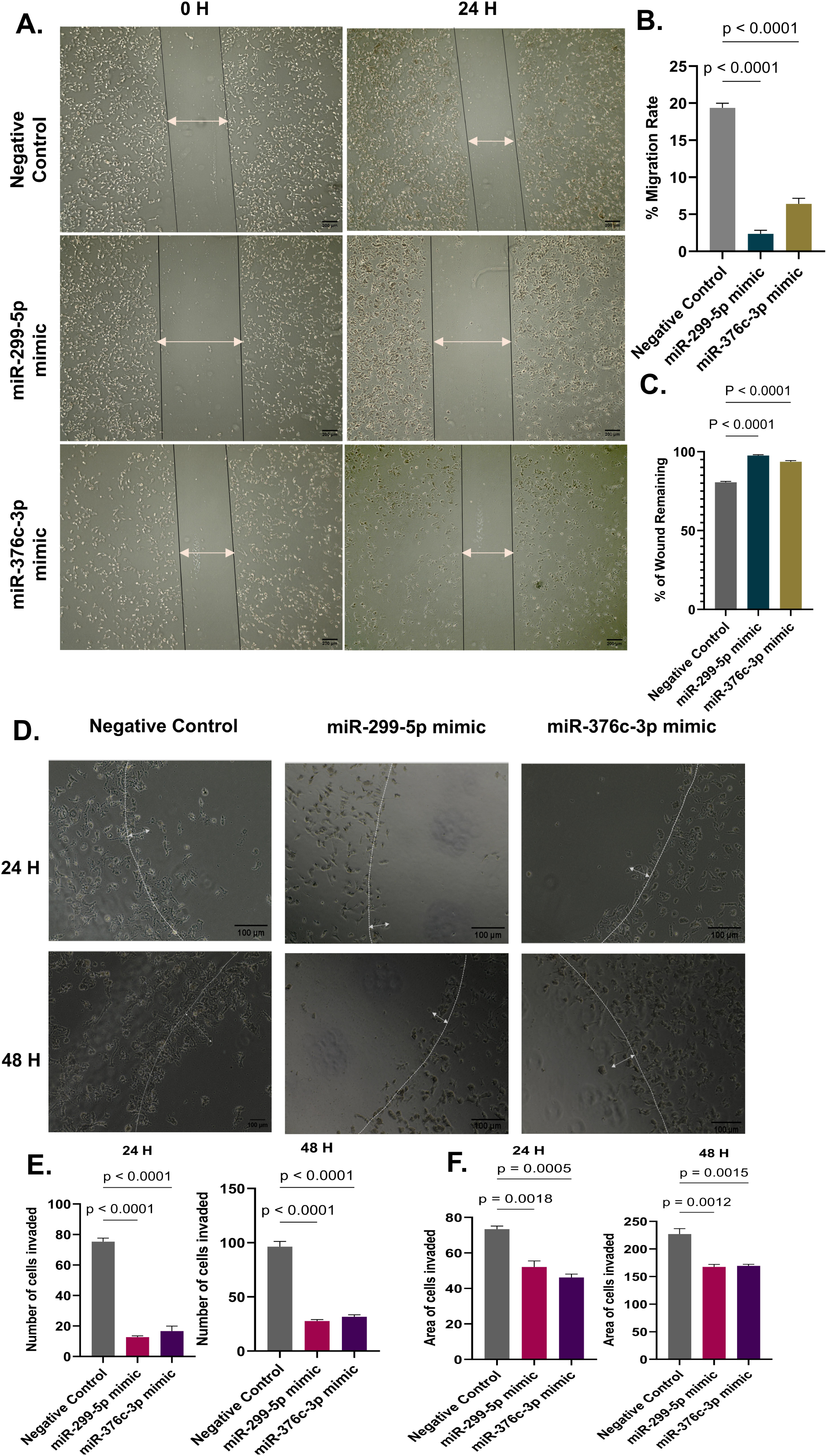
Restoring candidate C14MC miRNA expression inhibits Huh7 migration and invasion. (A) Micrographs showing inhibition of Huh7 migration and reduced wound healing upon transfecting with miR-299-5p or miR-376c-3p mimics compared to cells transfected with negative control. (B) Bar graph showing reduction in percentage migration of Huh7 cells upon mimics transfection compared to negative control (C) Bar graph showing percentage of wound remaining upon mimics transfection compared to the negative control. (D) Representative micrographs showing a reduction in invasion of Huh7 cells into the 2D-agarose spot post-transfection with miR-299-5p or miR-376c-3p mimics compared to cells transfected with negative control post 24H transfection. (E) Number of cells invading the 2D-agarose spot when transfected with miRNA mimics in comparison to cells transfected with negative control post 24H and 48H of transfection. (F) Area of the 2D-agarose spot invaded by when transfected with miRNA mimics in comparison to cells transfected with negative control post 24H and 48H of transfection.

### 2.7. miR-379/656 cluster and its target network are critical to predicting overall survival in HCC patients

We also assessed the clinical utility of C14MC candidate miRNAs and their targets, namely, miR-299-5p, miR-376c-3p, PARP1, SPP1, RAD21, and CENPA. KM survival analysis revealed that the overall survival of HCC patients was significantly correlated with SPP1 (**Figure 7A**) and CENPA expressions (**Figure 7B**). We observed that patients with lower expressions of SPP1 [HR=2.24, 95% CI (1.57-3.19), Log-rank p=5.2e-06] and CENPA [HR=2.3, 95% CI (1.63-3.26), Log-rank p=1.1e-06] had better median survival. Additionally, we checked if other co-founding clinical parameters, such as HCC stage, grade, patient gender, patient race, chemotherapy administration via sorafenib, alcohol consumption, and viral hepatitis (HBV), might affect the survival outcomes (**Table 1**). Furthermore, multivariate-Cox regression analysis was performed to assess the combined effect of C14MC and the validated gene targets. We observed that HCC stage and CENPA were significantly associated with all three survivals, overall survival (OS), disease-free survival (DFS), and progression-free survival (PFS). Other specific miRNA or target gene associations with individual survival outcomes showed SPP1 to be associated with OS; SPP1 and miR-376c-3p to be associated with DFS (**Figure 7C-7E**).

**Figure 7.**
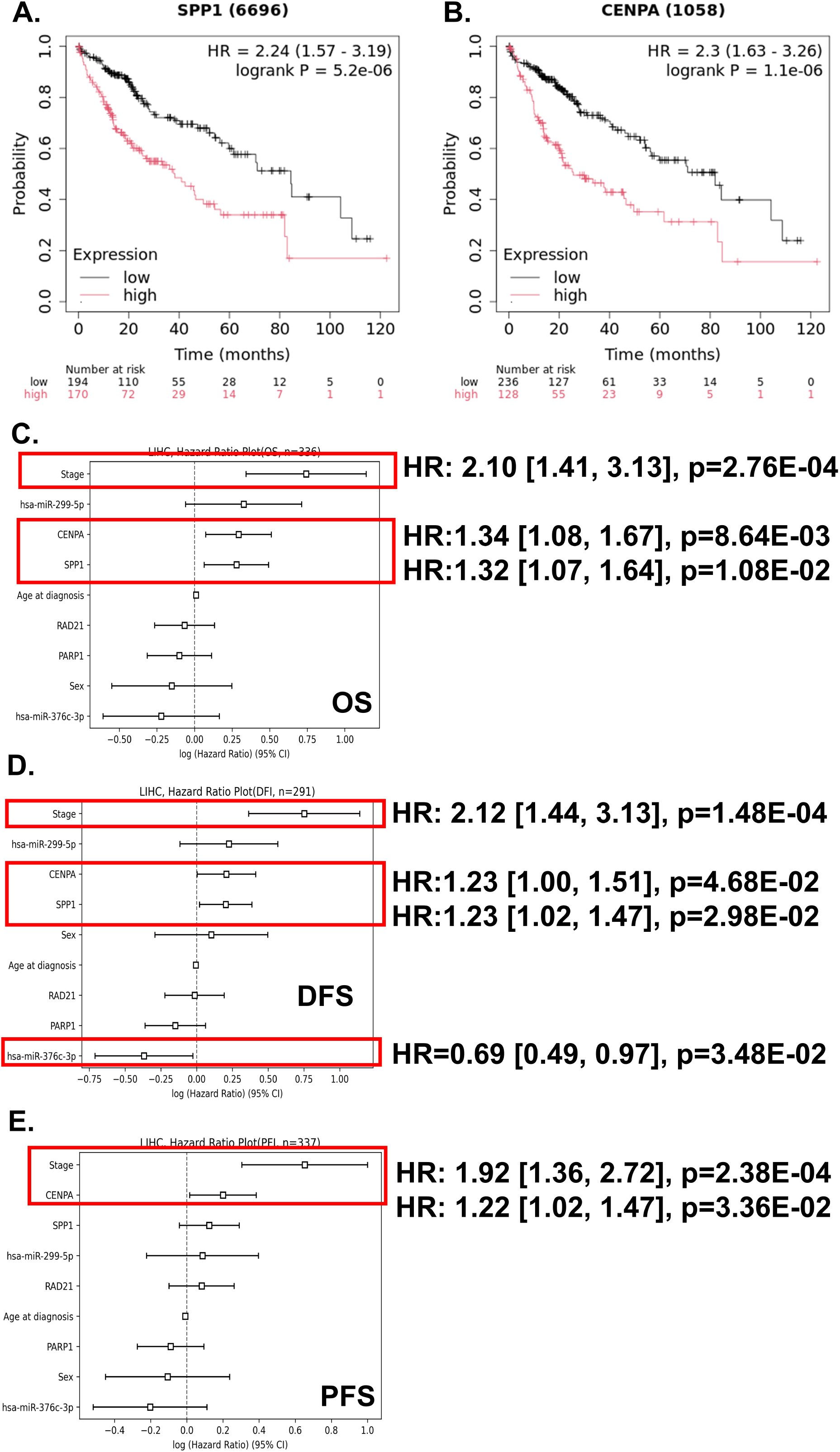
Clinical utility of C14MC miRNAs and gene targets in HCC. (A) and (B) KM plots of SPP1 and CENPA showing that these genes are significantly associated with the overall survival of HCC patients. Multivariate Cox regression analyses for miR-299-5p, miR-376c-3p, and their target genes PARP1, SPP1, CENPA, and RAD21, showing that (C) HCC stage, CENPA, and SPP1 expressions are significantly associated with overall survival, (D) HCC stage, CENPA, SPP1, and miR-376c-3p are significantly associated with disease-free survival, and (E) HCC stage and CENPA are significantly associated with progression-free survival, respectively.

**Table 1.** List of various factors that are significantly associated with overall survival based on C14MC target gene expressions among HCC individuals. (Survival analysis was performed using survival data and transcript expression data from the TCGA-LIHC cohort)

|  |  | Hazard Ratio (HR), 95% C.I. | Log-rank p value (**p<0.05**) |
| --- | --- | --- | --- |
| ***PARP1*** |  |  |  |
| Stage | 1 & 2 | 1.31 (0.77-2.22) | 0.31 |
|  | 3 & 4 | 2.51 (1.22-5.71) | 0.01 |
| Grade | 1 | 2.34 (0.76-7.14) | 0.13 |
|  | 2 | 1.83 (1.07-3.16) | **0.026** |
|  | 3 | 1.48 (0.78-2.80) | 0.23 |
| Gender | Male | 1.77 (1.08-2.92) | **0.023** |
|  | Female | 0.61 (0.88-2.93) | 0.099 |
| Race | White | 1.7 (1.03-2.8) | **0.035** |
|  | Asian | 1.61 (0.88-2.93) | 0.12 |
| Treatment | Sorafenib | 3.28 (0.71-15.2) | 0.11 |
| Alcohol consumption | Yes | 1.97 (1.03-3.8) | **0.038** |
|  | No | 0.73 (0.46-1.18) | 0.2 |
| Viral Hepatitis | HBV + | 1.97 (1.03-3.8) | **0.038** |
|  | HBV - | 1.56 (0.8-3.04) | 0.19 |
| ***SPP1*** |  |  |  |
| Stage | 1 & 2 | 2.31 (1.41-3.78) | **0.00059** |
|  | 3 & 4 | 2.15 (1.18-3.9) | **0.01** |
| Grade | 1 | 2.1 (0.81-5.45) | 0.12 |
|  | 2 | 2.07 (1.21-3.53) | **0.0066** |
|  | 3 | 3.44 (1.88-6.29) | **2e-05** |
| Gender | Male | 2.66 (1.69-4.17) | **1.1e-05** |
|  | Female | 1.99 (1.12-3.52) | **0.016** |
| Race | White | 1.9 (1.12-3.24) | **0.016** |
|  | Asian | 3.3 (1.78-6.11) | **5.7e-05** |
| Treatment | Sorafenib | 2.07 (0.69-6.21) | 0.19 |
| Alcohol consumption | Yes | 3.03 (1.38-6.63) | **0.0037** |
|  | No | 2.38 (1.27-4.44) | **0.0051** |
| Viral Hepatitis | HBV + | 3.16 (1.65-6.05) | **0.00024** |
|  | HBV - | 2.51 (1.54-4.08) | **0.00014** |
| ***RAD21*** |  |  |  |
| Stage | 1 & 2 | 0.82 (0.51-1.33) | 0.42 |
|  | 3 & 4 | 2.35 (1.28-4.34) | **0.0047** |
| Grade |  |  |  |
|  | 1 | 2.41 (0.92-6.28) | 0.065 |
|  | 2 | 1.28 (0.75-2.2) | 0.37 |
|  | 3 | 1.95 (1.04-3.65) | **0.034** |
| Gender | Male | 1.59 (0.98-2.57) | 0.059 |
|  | Female | 1.21 (0.69-2.1) | 0.5 |
| Race | White | 1.32 (0.83-2.11) | 0.24 |
|  | Asian | 2.35 (1.29-4.28) | **0.0038** |
| Treatment | Sorafenib | 2.29 (0.71-7.32) | 0.15 |
| Alcohol consumption | Yes | 1.52 (0.78-2.95) | 0.21 |
|  | No | 1.4 (0.84-2.36) | 0.2 |
| Viral Hepatitis | HBV + | 0.59 (0.3-1.19) | 0.14 |
|  | HBV - | 2.23 (1.34-3.7) | **0.0016** |
| **CENPA** |  |  |  |
| Stage | 1 & 2 | 2.55 (1.57-4.14) | **8.5e-07** |
|  | 3 & 4 | 2.18 (1.22-3.89) | **0.0069** |
| Grade | 1 | 4.75 (1.73-13.02) | **0.0011** |
|  | 2 | 2.22 (1.32-3.73) | **0.002** |
|  | 3 | 3.01 (1.61-5.63) | **0.00031** |
| Gender | Male | 3.94 (2.03-7.66) | **1.3e-05** |
|  | Female | 2.71 (1.48-4.96) | **0.00078** |
| Race | White | 1.8 (1.14-2.84) | **0.011** |
|  | Asian | 5.65 (3.02-10.57) | **1.3e-09** |
| Treatment | Sorafenib | 3.91×10<sup>8</sup> (0-∞) | **0.013** |
| Alcohol consumption | Yes | 2.95 (1.34-6.46) | **0.0048** |
|  | No | 2.65 (1.66-4.24) | **2.4e-05** |
| Viral Hepatitis | HBV + | 1.84 (0.95-3.54) | **0.066** |
|  | HBV - | 2.59 (1.64-4.1) | **2.6e-05** |

Stage 1 & 2 (n=257) 3 & 4 (n=90) Grade 1 (n=55) 2 (n=177) 3 (n=122)
Gender Male (n=250)
Female (n=121) Race White (n=194) Asian (n=158) Treatment Sorafenib (n=30) Alcohol consumption Yes (n=107) No (n=205) Viral hepatitis
HBV + (n=153) HBV – (n=169)

## 3. Discussion

Hepatocellular carcinoma (HCC) is the most common and lethal form of primary liver cancer (PLC). The current diagnostic HCC markers and treatment options available for HCC include a six-month ultrasound scan coupled with or without serum alpha-fetoprotein (AFP) measurements and sorafenib administration (35–37). Despite the improvements in terms of HCC screening, the overall survival of HCC patients remains poor (38,39). Novel transcriptomic markers have huge potential in terms of cancer diagnosis and therapy (40). C14MC or miR-379/656 cluster is the second-largest polycistronic miRNA cluster in humans that encodes nearly 50 different miRNAs (23). The aberrant expression of this cluster is implicated in the tumorigenesis of several cancers (28, 41,42). However, the functional regulation and role of this cluster in HCC pathophysiology have not been established. The present study aimed at understanding the mechanistic regulation of C14MC, especially through epigenetic mechanisms like DNA methylation, and its functions in HCC using *in vitro* models of HCC and various clinical datasets. This is the first study to delineate the regulatory mechanism exerted by DNA methylation on C14MC expression in HCC. We show that i. C14MC miRNAs are downregulated in HCC cells and tissues, suggesting a tumor-suppressor role of this cluster in HCC; ii. Promoter CpG hypermethylation suppresses C14MC in HCC cells and tissues; iii. C14MC miRNAs like miR-299-5p and miR-376c-3p regulate critical oncogene expression post-transcriptionally, and miRNA reactivation can abrogate PARP1, SPP1, RAD21, and CENPA oncogene expressions; iv. Reactivation of C14MC miRNAs directly inhibited cancer hallmarks such as cell migration and invasion; v. Several of these miRNAs and gene targets are critical to predicting the HCC patient survival outcomes. In summary, we report that C14MC downregulation is tightly regulated by DNA methylation, which drives the overexpression of several oncogenes targeted by C14MC. Thus, the C14MC methylome and its target transcriptome may be critical to HCC management, and designing novel RNA-based therapeutics can help mitigate HCC.

C14MC or miR-379/656 cluster is frequently dysregulated in various cancers. Depending on the specific cancer types, this cluster is known to have both oncogenic and tumor suppressor functions, which point towards diverse biological roles associated the cancer development and progression. C14MC downregulation is often reported in breast cancer, glioblastoma, oligodendroma, and cervical cancer (21,27,28,42). Contrastingly, oncogenic functions associated with the C14MC are identified in a few cancers (43). We showed that members of C14MC were significantly downregulated in HCC cell lines and tissue specimens from the LIHC cohort. This suggests that the cluster has a broader tumor suppressor function in HCC.

DNA methylation is perhaps the most significant epigenetic alteration that affects post-transcriptional gene regulation (22). Hypermethylation at upstream regulatory elements such as the promoter region can suppress downstream target genes (30). Abnormal promoter methylation might result in the development of various pathophysiological conditions beyond cancer (44). The hypermethylation at C14MC upstream regulatory elements is implicated in carcinoma of the cervix (54), glioblastomas (27), and oligodendromas (28). However, hypomethylation of the C14MC regulatory locus was reported in lung adenocarcinoma, Temple syndrome and conditions like atherosclerosis (44,45). We were interested in understanding the promoter methylation of C14MC in HCC cells and tissues. We identified the putative promoter and confirmed its activity by cloning it into a luciferase plasmid and subsequently performed the luciferase assay, which confirmed the promoter function associated with the genomic region. Further, we observed that upon artificial methylation, there was a substantial reduction in luciferase activity, which suggested that the promoter function is tightly linked to methylation levels and that methylation can reduce the C14MC promoter function. A mutual exclusiveness exists between promoter methylation and transcription factor binding, which can affect gene expression regulation (34). We identified through *in silico* analysis the different transcription factors that might bind to the C14MC. Notably, the HNF-4α binding site was identified within the C14MC promoter. Interestingly, HNF-4α is downregulated in HCC, and overexpressing HNF-4α can reverse HCC malignancy by regulating the genes associated with the DLK1-DIO3 locus, including the miRNAs belonging to this cluster, like miR-134, which are known to suppress the KRAS oncogene (46). Further, the data generated from bisulfite-Sanger sequencing and CpG methylome data of patient samples from different patient cohorts, including the TCGA-LIHC, suggested a prominent hypermethylation of specific CpG sites within the promoter of C14MC. These results point to an intricate regulatory mechanism associated with C14MC tumor suppressor function.

miRNAs are critical to regulating the post-transcriptional gene regulatory networks. Aberrant miRNA expressions can affect the downstream target gene expression (12). The involvement of such a dysregulated miRNA-mRNA axis is well studied in different cancers (18). Notably, C14MC dysregulation is shown to directly target PDK1, a critical gene in the PI3-Akt signaling in carcinoma of the cervix (42). We identified and validated the expression of post-transcriptional regulatory networks targeted by C14MC miRNAs in specific hub genes of HCC. PARP1 overexpression in HCC is often associated with resistance against sorafenib-induced cell death and radiotherapy (47–49). RAD21 is a reliable prognostic marker in HCC patients, with its expression generally associated with poorer differentiation status and tumor size (50–52). SPP1 overexpression in HCC is known to drive HCC cell proliferation and tumor growth (53), and higher SPP1 expression levels are associated with tumor macrophage infiltration and can determine survival outcome among HCC patients (54). We identified that PARP1, RAD21, and SPP1 as the direct targets of miR-299-5p of C14MC, and overexpressing miR-299-5p suppressed the expression of these oncogenes. Likewise, the expression of CENPA, a prominent cell cycle regulation gene with established roles in regulating key mitotic processes like mitotic protein assembly and chromosomal segregation (55,56), was abrogated in HCC cells upon miR-376c-3p overexpression in our study.

Cancer cells differ from non-cancerous cells in terms of their biological abilities (57). The emergence of altered cellular epigenetic reprogramming in the plasticity of cancerous cells is considered a hallmark of cancer (58). We checked the ability of C14MC restoration to inhibit HCC cell proliferation and invasiveness through cellular assays. Dysregulated cell proliferation may promote HCC advancement and progression (59). The ectopic expression of C14MC miRNAs-miR-299-5p and miR-376c-3p, in HCC cells, inhibited HCC cell migration and invasion. Previously, it was reported that miR-299-5p suppresses the migratory ability and invasiveness of papillary thyroid carcinoma (60) and breast cancer (61). Further, overexpressing another C14MC member, miR-376c-3p, is known to inhibit cancer cell proliferation, migration, and invasion in oral squamous cell carcinoma (OSCC) (62), malignant gastric cancer (63), and medullary thyroid carcinoma cells (64). Furthermore, our pathway enrichment of the hub genes identified that targets of C14MC, like PARP1 and SERPINE1, in regulating apoptosis and cellular senescence, which are critical to determining the cellular proliferation, migration, and invasion capabilities. Thus, we show that reactivation of C14MC or its component miRNAs may inhibit cell migration and invasion and thereby might contribute to mitigating HCC progression.

Lastly, we identified the clinical significance of C14MC in HCC. Several studies have shown that members of C14MC can be useful as diagnostic and prognostic markers in various cancers (12). However, the clinical utility of the region in HCC remains elusive. We have previously shown that individual miRNAs of this cluster can be used as a potential diagnostic marker for HCC (16). In this study, we report the use of two panels of C14MC miRNAs-miR-382-5p and miR-376c-3p, and miR-299-5p, miR-376c-3p, miR-656-3p, miR-376a-3p and miR-377-3p, which can be used as potential diagnostic panels for HCC. Additionally, we have shown that the C14MC members and their gene targets can be employed to predict survival outcomes in HCC patients (OS, DFS, and PFS). Interestingly, we observed that the expressions of several of these targets were significantly associated with gender, race, HCC grade, cancer stage, treatment strategies, and other HCC accompanying risk factors such as alcohol consumption and prevalence of viral hepatitis. Taken together, the observations show that C14MC and its target gene expressions might not just help predict survival outcomes in HCC patients, but also might help design strategies that can help improve the survivability of the patients, depending on the clinical parameters of the disease, like histopathological outcomes and risk factors associated with HCC.

## 4. Conclusions

We showed that members of C14MC are downregulated in HCC and that the miRNA cluster acts as a tumor suppressor in HCC. We also showed that the underlying epigenetic cause behind the loss of tumor suppressor function in HCC is promoter hypermethylation. A systems biology approach was used to identify the C14MC transcriptome, which was subsequently validated experimentally. Additionally, the key ontologies and pathways regulated by the transcriptomic network were identified, and the key cancer hallmarks of HCC, including cell migration and invasion upon cluster miRNA activation, were experimentally shown. We also showed that downregulation of the cluster is significantly correlated with disease prognosis and can be useful to develop strategies in clinical settings for HCC management. Although we discuss the role of methylation in C14MC regulation and functions of some of the miRNAs of C14MC in this study, additional studies targeting the entire C14MC and establishing C14MC-activated HCC *in vitro and in vivo* models can help characterize the actual transforming ability of the cluster and study its biological functions during HCC carcinogenesis.

## 5. Methods and Methods

### 5.1. Cell culture and Spheroid Culture

We procured HCC cell lines from NCCS, Pune, India, through their standard procurement procedure (http://ncmr.nccs.res.in/home). The cell lines (HepG2: MEM, and Huh7: DMEM: F12 Ham’s) were maintained with 10% fetal bovine serum supplementation. For spheroid generation, HepG2 cells were used, and the tumuroids were generated by using a modified forced suspension method as described previously (65). The control RNA samples were procured from TakaraBio, Japan.

### 5.2. Promoter identification, cloning, characterization

We identified the putative promoter of C14MC using the FANTOM5 tool (66). The promoter region was then cloned into the pGL3-Basic plasmid between XhoI and HindIII restriction sites to generate the pB-C14MC construct. The successful cloning of the promoter region into the pGL3-Basic vector was confirmed by double digesting the construct with XhoI and HindIII and diagnostic digestion with KpnI restriction enzymes. We then performed artificial methylation experiments using the pB-C14MC construct by incubating it with M.SssI methyltransferase enzyme (ThermoFisher Scientific, USA). The artificially methylated and unmethylated constructs were then co-transfected with the SV40 plasmid using Fugene HD Reagent (Promega, USA). The reporter activity was then measured using a dual luciferase assay kit (Promega, USA) 48h post-transfection using a GloMax 20/20 Luminometer.

### 5.3. C14MC promoter partial methylation status identification in HCC cells

The partial methylation status of the identified promoter in HCC cells was assessed by performing bisulfite PCR, followed by sequencing the PCR products by Sanger sequencing. The MethPrimer tool was used to design the bisulfite-specific PCR primers (67). To determine the methylation status, the genomic DNA was isolated from HCC cells and modified using the EZ DNA Lightening Conversion Kit (Zymo Research, USA). The converted DNA was then amplified using the primers FP: GTTTATATTTGGGAATTAGTTATGT and RP: TCAAACACAATATATAAAAAAAATC in a thermocycler (Applied Biosystems, USA). The amplification conditions included: 95°C for 5 min, 34 cycles at 94°C for 90 sec, 54.5°C for 3 min, 72°C for 1 min, and 1 cycle at 72°C for 5 min. The PCR products were then gel-purified using the E.Z.N.A. Gel Extraction kit (Omega Bio-tek, Georgia). The sequence trace files were then analysed using the BiQ Analyzer software to calculate the percent methylation at individual CpGs (68).

### 5.4. qRT-PCR

We performed the first strand cDNA synthesis of mRNA using the Verso cDNA Synthesis kit (ThermoFischer Scientific, USA). For miRNA expression analysis, cDNA conversion was performed using the miRCURY LNA RT kit (Qiagen, Germany). qRT-PCR for miR-299-5p, miR-376c-3p, miR-377-3p, miR-376a-3p, and miR-656-3p were performed using the miRCURY LNA SYBR Green PCR kit (Qiagen, Germany). qRT-PCR for PARP1, SPP1, RAD21, and CENPA were performed using the SYBR DyNAmo ColorFlash SYBR Green (ThermoFischer, USA). Β-Actin and U6 snRNA were used as internal controls. The experiments were performed using a Rotor-GeneQ system (Qiagen, Germany). The relative fold change in the expression was calculated using the 2^-ΔΔCT^ method. The primer details used in this study are provided in **Supplementary Table 1.**

### 5.5. NanoString nCounter miRNA expression analysis

We performed the nCounter Human v3 miRNA Expression Assay (NS_H_miR_v3B) on an nCounter Analysis System (NanoString Technologies) according to the guidelines by the manufacturer to identify the differentially expressed miRNAs (69). We manually screened for differentially expressed C14MC miRNAs out of the 799 endogenous miRNAs present in the panel. The normalization was performed using geometric means of positive controls and the top 100 highly expressed miRNAs (−1.5≤log_2_(FC)≥+1.5, p≤0.05).

### 5.6. Whole Transcriptomic analysis

RNA-seq was performed to identify the expression of the C14MC target genes in HCC tumouroids. We performed RNA-seq on an Illumina NovaSeq V1.5 instrument according to the manufacturer’s protocol. Briefly, the ribosomal RNA depletion was carried out using Qiagen FastSelect rRNAHMR (Qiagen, Germany), and the mRNA libraries were prepared using the NEB Ultra II directional RNA-Seq Library Kit (NEB, USA). The first-strand and second-strand cDNA synthesis were carried out using reverse transcriptase and DNA Polymerase I enzymes, respectively, followed by adapter ligation. Prior to sequencing, the fragment sizes were analysed using HS NGS Fragment kit (1-6000bp) (Agilent, USA) on a Fragment analyser. Subsequently, the mapping of the aligned paired reads was performed against the reference genome (GRCh38) using STAR (v.2.7.11b). The DETGs were identified using DESeq2 (70). The fold change was calculated using the thresholds (−2≤log_2_(FC)≥+2, p≤ 0.05).

### 5.7. TCGA-LIHC cohort analysis

We analysed the expressions of C14MC in The Cancer Genome Atlas-Liver Hepatocellular Carcinoma (TCGA-LIHC) as described previously in our study (16). Further, the diagnostic potential of C14MC expression was identified by calculating the sensitivity and specificity by plotting ROC curves using the sum of expressions of individual miRNAs. The target genes of C14MC were identified using the MIENTURNET tool by querying the individual miRNAs in the miRTarBase (71). The prognostic significance of C14MC and its target genes in LIHC was analysed using the DoSurvive tool (72). The protein-protein interaction (PPIN) network analysis was constructed using the STRING web-based tool. Further, the top ten hub gene networks were predicted using the MCC algorithm using the CytoHubba tool (73). Furthermore, the gene ontology associations in terms of biological processes (BPs), molecular functions (MFs), pathway analyses, and disease associations were identified using the ShinyGO online tool (74). Further, the CpG methylation status in the TCGA-LIHC cohort and other independent datasets were analysed using the Epigenome-Wide Association Study (EWAS) data hub (75).

### 5.8. Migration assay

We transfected Huh7 cells with miR-299-5p or miR-376c-3p mimics or All-Stars-Negative Control (Qiagen, Germany) according to the recommendations from the manufacturer using the TrasIT-X2 Dynamic Delivery System (Mirus Bio, USA). Before transfection, the cells were subjected to serum starvation for 24h. A wound was made in the center of the plate using a sterile micropipette tip. Post-transfection, complete medium containing 10% FBS was supplemented, and the cell migration into the wounded region was monitored using an EVOS XL Core imaging system (ThermoFischer Scientific, USA).

### 5.9. 2D-agarose spot invasion assay

We used a modified protocol of the 2D-agarose spot invasion assay from what was described earlier (76). Briefly, Huh7 (3×10^5^ cells) were seeded using serum-free DMEM: F12 Ham’s into P35 cell culture dishes containing 0.5% agarose spots with 20% FBS. Post 4h of incubation, the media was removed and the cells were transfected with miRNA mimics (miR-299-5p or miR-376c-3p) or negative control. The cells that invaded the agarose spots were imaged at 10X magnification and were scored from at least three microscopic fields manually to determine the extent of cell invasion.

### 5.10. Statistical analysis

The statistical significance was computed either by a Student’s *t-*test or One-way ANOVA, and a p-value ≤0.05 was considered to be statistically significant. All the data are represented as means±Standard Error Means (SEMs). All the experiments were performed in biological replicates.

## Data availability

The data discussed in this study are enclosed in the article or are provided in the supplementary information. The data supporting the findings discussed in the article is available upon a suitable request made to the corresponding author.

## Author contributions: CRediT

Shreyas Hulusemane Karunakara: Writing-original draft, Conceptualization, Methodology, Investigation, Validation, Formal analysis, Funding acquisition. Gopalakrishna Ramaswamy: Methodology, Data curation. Shama Prasada Kabekkodu: Writing-Review and editing, Conceptualization, Investigation, Validation, Supervision. Akila Prashant: Writing-Review & editing, Supervision. Prashant Vishwanath: Writing-Review & editing, Resources. Rohit Mehtani: Writing-Review & editing, Prasanna Santhekadur: Writing-Review & editing, Conceptualization, Supervision, Funding acquisition.

## Ethics Declaration

None

---

**Supplementary Table 1.**
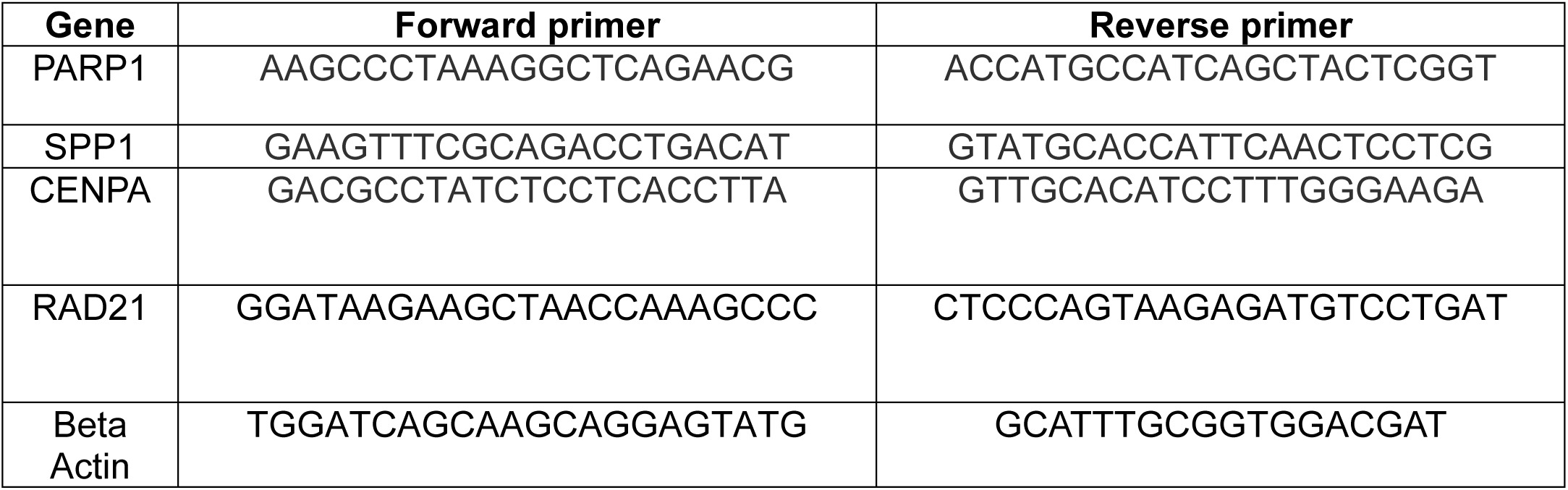
Primer sequences used for qRT-PCR expression studies.

